# Multiplexed Gene Synthesis in Emulsions for Exploring Protein Functional Landscapes

**DOI:** 10.1101/163550

**Authors:** Calin Plesa, Angus M. Sidore, Nathan B. Lubock, Di Zhang, Sriram Kosuri

## Abstract

Next-generation sequencing has engendered an expanding suite of functional assays that can test sequence-function relationships at unprecedented scales in pooled formats (multiplex). Such assays are currently constrained by the short length of oligonucleotide (oligo) pools, which limit potential applications. Here we report a simple, low-cost, and scalable method called DropSynth that assembles gene libraries from oligo pools for use in multiplexed functional assays. DropSynth utilizes a library of barcoded beads to isolate and concentrate oligos needed for a gene’s synthesis in a pooled format. These bead-bound oligos are then emulsified, processed, and assembled into genes within the emulsion droplets. We synthesized ~1000 phylogenetically diverse orthologs of phosphopantetheine adenylyltransferase (PPAT) and tested their fitness in a multiplexed functional assay. While the majority of orthologs complement, those that do not are broadly distributed across the phylogenetic tree. Synthetic errors in our assemblies allow us to explore local landscapes around the designed orthologs revealing constrained mutations for complementing orthologs as well as gain-of-function mutations for low-fitness orthologs. This broad mutational scanning approach is complementary to deep mutational scanning and helps us understand proteins by probing evolutionarily divergent sequences that share function.

---

The rapid progress in DNA sequencing has provided a glimpse of the vast diversity of sequence and function that nature has explored. Unfortunately our ability to explore the functional aspects of diverse sequences remains minuscule. Multiplexed functional assays are a new set of techniques (also called Massively Parallel Reporter Assays^1^ and Multiplexed Assays for Variant Effects^2^) that link a function to a next generation sequencing (NGS)-based output to allow testing of thousands to millions of sequences for function in a pooled manner. Mutagenesis and deep mutational scanning combined with multiplexed functional assays are now able to systematically probe all single and double amino acid substitutions for a protein’s function^3,4^. Despite the power of these approaches, the sequence space explored in such experiments is minute when compared to the evolutionary distance between even highly-homologous protein sequences. Our ability to sample a broad diversity of sequences is bottlenecked by two limitations in existing approaches for producing the libraries to be tested. First, low-cost microarray-based oligo libraries allow for large libraries of designed <200 nucleotide (nt) sequences^5^, which is far below the typical length of a protein and limits many other applications. Secondly, gene synthesis is capable of creating long-length sequences, but its costs currently make it prohibitively expensive for sampling large libraries of designed sequences^6–9^.

Here we develop a new gene synthesis method we term DropSynth, a multiplexed approach capable of exploring functional landscapes of gene-length sequences at much higher throughputs than currently possible. DropSynth uses low-cost oligo libraries and assembles gene libraries that are easily coupled into multiplex functional assays. We and others have developed robust parallel processes to build genes from oligo arrays, but because each gene must be assembled individually, costs are prohibitive for large gene libraries^6,10^. In these efforts, the ability to isolate and concentrate DNA from the background pool complexity was paramount for robust assemblies^11^. Previous efforts to multiplex such assemblies have not isolated reactions from one another, and thus suffered from short assembly lengths, highly-biased libraries, the inability to scale, and constraints on sequence homology^12–15^.

DropSynth works by pulling down only those oligos required for a particular gene’s assembly onto barcoded microbeads from a complex oligo pool. By emulsifying this mixture into picoliter droplets, we isolate and concentrate the oligos prior to gene assembly thereby overcoming the critical roadblocks for proper assembly and scalability (Fig. 1A). The DropSynth barcodes are unique 12 nt sequences that all oligos for a particular assembly share, and pair with complementary strands displayed on the microbead. Within each droplet, sequences are released from the bead using Type IIs restriction enzyme sites and assembled through polymerase cycling assembly (PCA) into full length genes. Finally, the emulsion is broken and the gene library is recovered. To test and optimize the protocol, we built model assemblies that were unique, but shared common overlap sequences. As a result, any contaminating oligo would still participate in the assembly reaction, allowing us to monitor assembly specificity and library coverage. We optimized each aspect of the protocol by trying to assemble 24-, 96-, and 288-member libraries composed of 3, 4, 5, and 6 oligos at once based on how often we saw intended targets versus their expected frequency given random (i.e. bulk) assembly (Fig. 1B). Over many iterations, we achieved high enrichment rates (~10^8^). Using these protocols, we were able to build libraries of up to 6 oligos that produced correct sized bands (Fig. 1C), and the resulting assembly distributions were not overly skewed (Fig. 1D).

**Figure 1.**
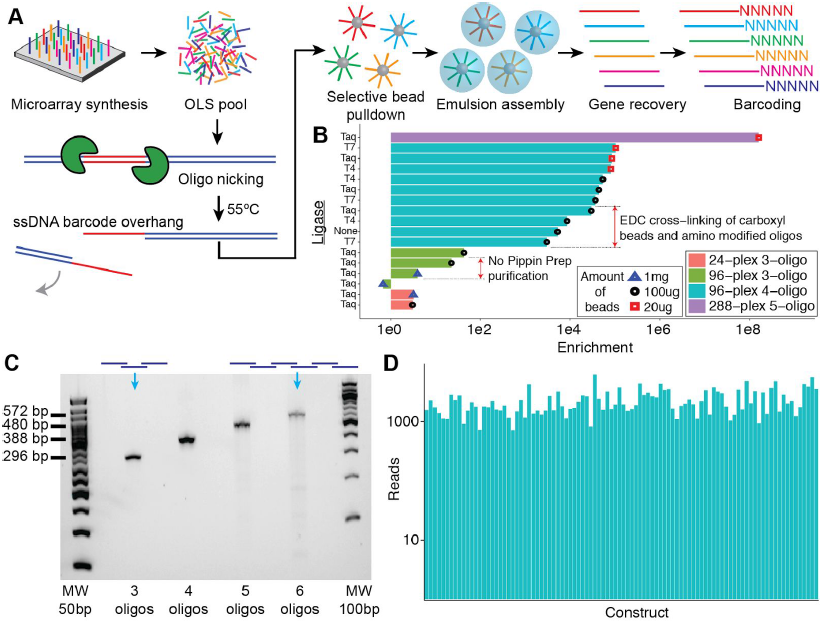
DropSynth assembly and optimization. **A)** We amplified array-derived oligos and used a nickase to expose a single-stranded overhang region that acts as a gene-specific barcode. Barcoded beads display complementary single-stranded regions that selectively pull down the oligos necessary to assemble each gene. After pulldown, we emulsified the beads and into droplets where a Type IIs restriction enzyme removes the barcodes and the liberated fragments are assembled by PCA. The emulsion is then broken, and the resultant assembled genes are barcoded and cloned. **B)** We used a model gene library that allowed us to monitor the level of specificity and coverage of the assembly process. We then optimized various aspects of the protocol including purification steps, DNA ligase, and bead couplings to improve the specificity of the assembly reaction. Enrichment is defined as the number of specific assemblies observed relative to what would be observed by random chance in a full combinatorial assembly. **C)** We attempted 96-plex gene assemblies with 3, 4, 5, or 6 oligonucleotides and ran the resultant libraries on an agarose gel. **D)** The relative read-counts of all 96 assemblies (4-oligo assembly) as determined by NGS were within the same order of magnitude.

As a proof-of-concept, we used DropSynth to assemble 1,152 orthologs of the *E. coli* enzyme phosphopantetheine adenylyltransferase (PPAT) from across the tree of life (Supp. Fig. 1). Each gene was assembled from five oligos. The median gene-length of the library was 483 basepair (bp) (range: 381-516 bp), and the median amino acid identity was 50% (Supp. Fig. 2A, B). We improved the assembly process to carry out the oligo isolation using a mixture of 384 uniquely barcoded beads (DropSynth barcodes). An oligo library synthesis (OLS) pool of 5,880 oligos of 200 nt, including three batches in 384-plex was synthesized on an Agilent microarray, assembled, and linked to random 20 bp barcode sequences (assembly barcode). We obtained correct-length bands for all three emulsion assemblies, in contrast to attempts using bulk assembly which produced shorter failed by-products (Supp. Fig. 2C, D). Sequencing revealed that a total of 872 genes (75%) had assemblies corresponding to a perfect amino acid sequence represented by at least one assembly barcode, with a median of 2 reads per assembly barcode and 56 assembly barcodes per ortholog (Fig. 2B, Supp. Fig. 3A, B). This coverage increased when including sequences with deviations from the designed sequences, with 1,002 genes (87%) represented within 5 aa from the designed sequences (all orthologs have some alignments regardless of distance). Among assemblies with at least 100 BCs, we observed a median of 1.9% perfect protein assemblies (*σ* = 2.9%) (Supp. Fig. 3C). The resultant error profiles were consistent with Taq-derived mismatch and assembly errors that we have observed previously^16^ (Supp. Fig. 4A, B). This suggests future improvements in polymerase optimization, oligo purity, and improved design algorithms will improve yield substantially.

**Figure 2.**
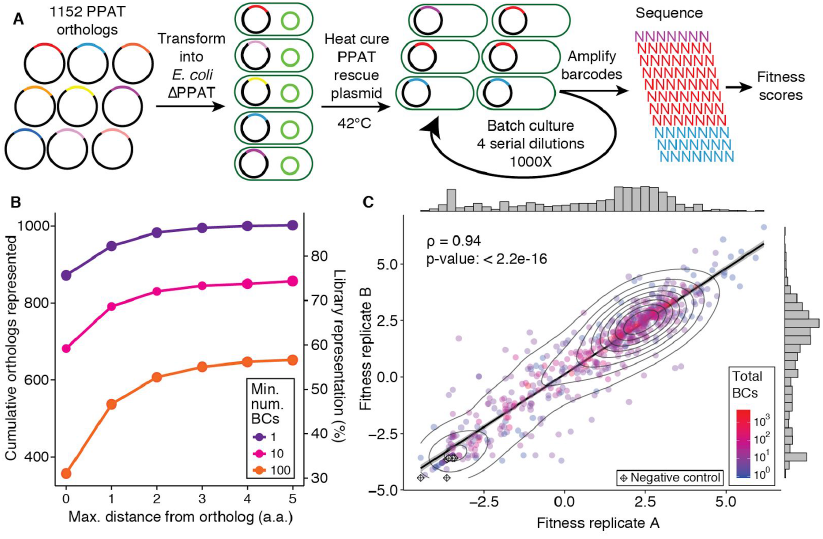
PPAT complementation assay. **A)** We used DropSynth to assemble a library of 1152 orthologs of phosphopantetheine adenylyltransferase (PPAT), an essential enzyme catalyzing the second-to-last step in coenzyme A biosynthesis, and tested using a pooled complementation assay. The barcoded library was transformed into *E. coli ΔcoaD* cells containing a curable rescue plasmid expressing *E. coli coaD*. The rescue plasmid was removed allowing the orthologs and their mutants to compete with each other in a batch culture. We tracked barcode frequencies over four serial 1000-fold dilutions, and used the frequency changes to assign a fitness score. **B)** We observe that 872 orthologs (75%) had at least one perfect assembly, and we observed 1,002 orthologs (87%) having at least one assembly within a distance of 5 a.a. from design. **C)** The fitness values for 651 orthologs across two independent biological replicates shows strong correlation (ρ=0.94; Pearson). Six negative controls lacking the H/TxGH motif required for nucleophilic attack on the α phosphate of the ATP have very low fitness values (<3) in the assay. We find that reproducibility among replicates improves with increasing number of barcodes (Supp. Fig. 7B).

**Figure 3.**
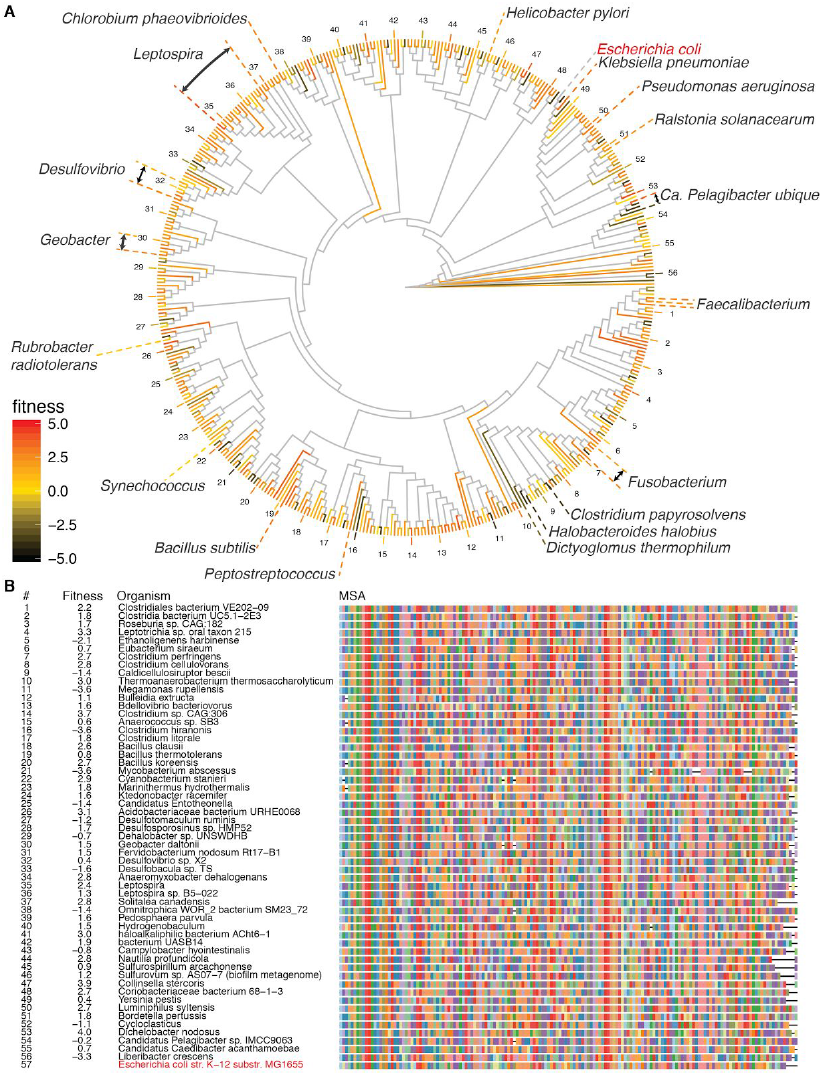
Complementation analysis of diverse orthologs **A)** We find that the majority of PPAT orthologs are able to complement, with 70% having positive fitness values, while low-fitness orthologs are dispersed throughout the tree without much clustering of clades. This phylogenetic tree shows 451 orthologs each with at least 5 barcodes, a subset of the full data set, where leaves are colored by fitness. **B)** Despite having a median 50% sequence identity, distant orthologs are typically still able to complement the function of the native *E. coli* PPAT (bottom row). This multiple sequence alignment shows 57 diverse PPAT orthologs, with highly gapped positions removed, along with their corresponding source organism and fitness scores.

**Figure 4.**
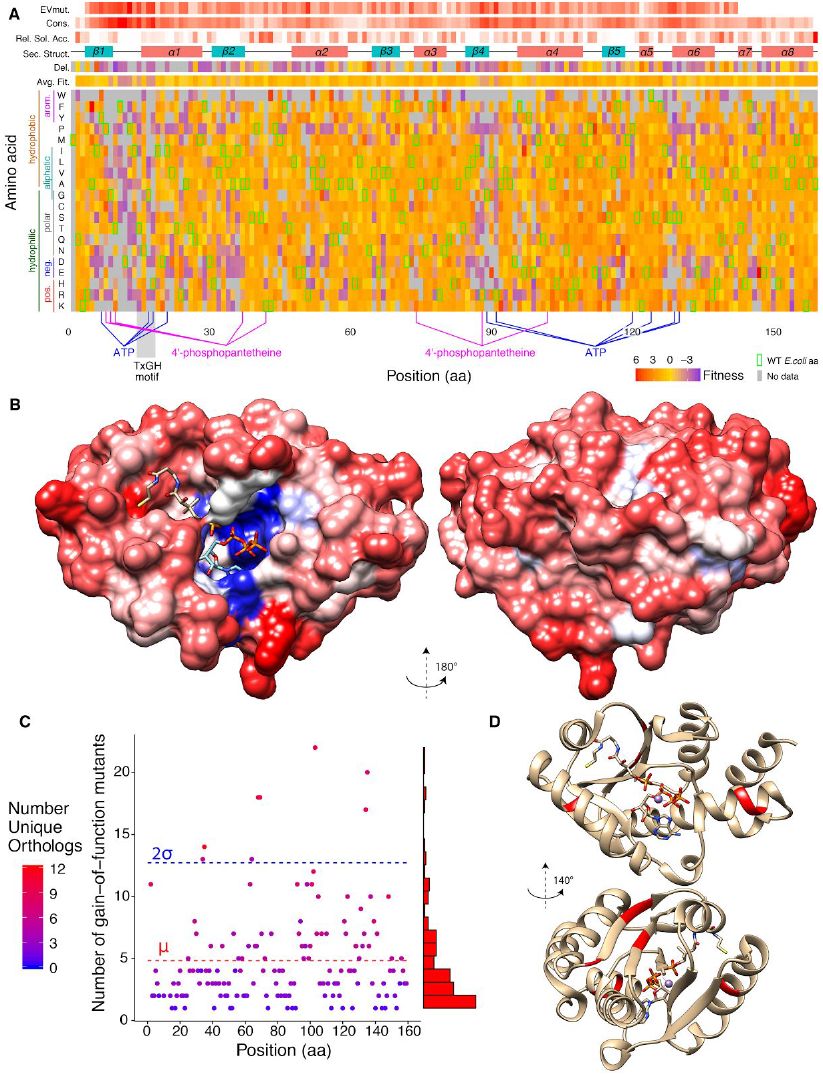
Broad mutational scanning (BMS) analysis **A)** A representation of the background-averaged sequence-function landscape of PPAT projected onto the *E. coli* PPAT sequence. Mutations are highly constrained at a core group of residues involved in catalytic function. Other positions show relatively little loss of function, when averaged over many orthologs, despite known interactions with the substrates. The matrix contains the mean fitness value derived from all 497 complementing orthologs and their 71,061 mutants within a distance of 5 a.a., at each aligned position. The *E. coli* WT sequence is indicated by green squares, while the average position fitness, fitness of a residue deletion, mean EVmutation evolutionary statistical energy^35^, site conservation, relative solvent accessibility, and secondary structure information is shown above. **B)** The *E. coli* PPAT (PDB: 1QJC, 1GN8^36^) structure complexed with 4’-phosphopantetheine and ATP shows the loss-of-function for mutations occurring at the active site, while other residues involved with allosteric regulation by coenzyme A or dimer interfaces show large promiscuity highlighting different strategies employed among orthologs. **C)** In addition to complementing orthologs we can also analyze mutants of low-fitness orthologs for gain-of-function (GoF) mutantions. Of the 129 orthologs with fitness less than -2.5, we found 385 GoF mutants with positive fitness for 55 orthologs. The position of the GoF mutations aligned onto the *E. coli* PPAT sequence, shaded by the number of orthologs which had at least one GoF mutation at this position. A total of 8 statistically significant positions (34, 35, 64, 68, 69, 103, 134, 135) were found corresponding to four regions in PPAT. **D)** *E. coli* PPAT structure with GoF residues shaded in red. Glu-134 is involved in hydrophobic interactions with coenzyme A^24^, suggesting a role for GoF mutations in modulating the inhibitory feedback, while Ala-103 participates in hydrophobic interactions between the PPAT dimers^17^.

We probed how well orthologs of various evolutionary distance to *E. coli* could rescue a knockout phenotype. PPAT is an essential enzyme, encoded on gene *coaD*, catalyzing the 2^nd^ to last step in the biosynthesis of coenzyme A (CoA) by transferring an adenylyl group from ATP to 4’-phosphopantetheine^17^ (Supp. Fig. 5). PPAT lacks homology to mammalian counterparts making it an attractive target for the development of novel antibiotics^18^. We created *E. coli ΔcoaD* cells with a heat curable rescue plasmid containing constitutively expressed wild-type *coaD* (Supp. Fig. 6A, C, D). This rescue plasmid showed escape frequencies lower than 1 in 20,000 under heat selection in pooled assays (Supp. Fig. 6E). Assembled PPAT variants on a barcoded high-copy plasmid (Supp. Fig. 6B) were transformed into *E. coli ΔcoaD* cells and screened for complementation by growing the library through in batch culture through three serial 1000-fold dilutions (Fig. 2A, Supp. Table 1). Barcode sequencing of the resulting populations provided an estimate for the fitness of all orthologs and mutants present. A fitness value of zero represents the growth rate necessary to maintain a stable population in the competitive assay, while large negative fitness values correspond to low growth rates leading to population depletion. Two biological replicates show strong correlation in ortholog fitness values (⍴=0.94; Pearson) which increases with the number of barcodes (Fig. 2C, Supp. Fig. 7A, B).Several negative controls, consisting of N-terminal truncated orthologs missing the H/TxGH motif required for catalytic activity, show strong depletion (Fig. 2C, Supp. Fig. 8A). Similarly, sequences containing indels leading to low sequence identities tend to quickly drop out of the population, and fitness is reduced with increasing numbers of mutations (Supp. Fig. 8B, C, 9). Fitness values are relatively noisy, with typical standard deviations of around 2.4 (Supp. Fig. 8D). We validated our fitness scores by using primers corresponding to the barcode to retrieve and individually test a subset of orthologs (Supp. Fig. 10). Sequence-verified clones were obtained for 37 of 49 orthologs and grown in a plate reader. The measured growth rates correlated well with the pooled fitness score (r_s_=0.86, Spearman) (Supp. Fig. 10B). The residual error of the resulting fit inversely correlated with the number of barcodes, highlighting the need for many barcodes to accurately determine fitness (Supp. Fig. 10C). Approximately 14% percent of the orthologs show strong depletion (fitness below -2.5) while 70% have a positive fitness value in the pooled assay. Low-fitness orthologs are evenly distributed throughout the phylogenetic tree with only minor clustering of clades (Fig. 3A, Supp. Fig. 11, 12A).

There are several reasons orthologs could have low fitness including environmental mismatches, improper folding, mismatched metabolic flux, interactions with other cytosolic components, or gene dosage toxicity effects resulting from improperly high expression. Of the orthologs from extremophilic bacteria, only alkaliphiles showed a slight propensity towards lower fitness values (Supp. Fig. 12B). Metabolic mismatch is unlikely since so many orthologs were able to complement well and both CoA and dephospho-CoA act as inhibitors implementing negative feedback loops to control the metabolic flux through the pathway^17^. Control experiments revealed that high expression levels of wild-type *E. coli* PPAT result in growth defects, while similar levels of expression for many other orthologs had no impact (Supp. Table 2). This observation parallels similar findings for *E. coli* DHFR where wild-type overexpression was toxic while overexpression of orthologs had no detrimental effects^19^, an effect linked to evolved protein-protein interactions that confer benefits at physiological concentrations. PPAT interaction partners include several enzymes encoded by essential genes such as *leuS*, *murE,* and *rplD*^20,21^.

Errors during the oligo synthesis or DropSynth assembly give us mutational data across all the orthologs, which we can further analyze to better understand function. We selected all 497 orthologs which showed some degree of complementation (fitness greater than -1) as well as their 71,061 mapped mutants within distance 5 a.a. and carried out a multiple sequence alignment to find equivalent residue positions. For each amino acid and position, we found the mean fitness among all of these orthologs and mutants. The resulting data was projected onto the *E. coli* PPAT sequence, as shown in Fig. 4A. This approach is akin to deep mutational scanning, except that the resultant data is an average fitness across all of the orthologs, rather than a particular sequence, in a manner more reminiscent of recent bioinformatic statistical coupling methods^22,23^. We term this approach broad mutational scanning (BMS). The average BMS fitness for each position shows strong constraints in the catalytic site and as expected, fitness is constrained at highly conserved sites and at buried residues compared to solvent-accessible ones (Supp. Fig. 13A, 13B). One surprising observation is that some residues which are known to interact with either ATP or 4’-phosphopantetheine turn out to be relatively promiscuous when averaged over a large number of orthologs. Furthermore, when mapped onto the *E. coli* structure (Fig. 4B), positions known to be involved with allosteric regulation by coenzyme A or dimer formation, show relatively little loss of function, highlighting the diversity of distinct approaches employed among different orthologs, while maintaining the same core function. Although 87% of the 3,180 possible mutations are covered, the coverage is strongly correlated with position fitness (Supp. Fig. 13C), implying that many mutations which are depleted in the pooled assay (and typically represented by a only a few barcodes), never pass the 10-read threshold used to filter assembly barcodes, an issue which can be resolved by sequencing the initial library to a greater depth. Unlike traditional mutagenesis approaches, the presence of multi-bp deletions from the oligo synthesis process also allows us to evaluate the effect of removal of entire residues from the sequence (Del. in Fig. 4A). Additionally, we can use those orthologs that did not complement and search for gain-of-function (GoF) mutations. A total of 385 gain-of-function (GoF) mutants out of 4,658 were found for 55 orthologs out of 129 low-fitness orthologs. By aligning these mutations to the *E. coli* sequence, the eight statistically significant residues (34, 35, 64, 68, 69, 103, 134, 135) shown in Fig. 4C localize to four small regions in the protein structure (Fig. 4D). In *E. coli*, residue Glu-134 and proximal Leu-102 have hydrophobic interactions with the cysteamine moiety of CoA^24^, suggesting some GoF mutations’ role in tuning CoA inhibition, while Ala-103 participates in hydrophobic interactions contributing to dimer formation^17^. Residues 64, 68, 69 are surface-exposed in the hexameric PPAT complex and are possible candidates for interactions with other proteins. As many of these mutations had only a single barcode, we estimated a false positive rate of 0.9% derived from the number of positive fitness mutants for negative controls (Supp. Fig. 8A). We retrieved six GoF mutants of six different orthologs from the library, each with fitness determined from only a single barcode, and individually tested their growth rates. Five of the six mutants showed strong growth and one failed to complement (Supp. Fig. 10B). We also tested two of the corresponding low-fitness orthologs, finding increases in the growth rate of 10% and 42% for their GoF mutants (Supp. Table 2).

Broad mutational scanning using DropSynth is a new tool to explore protein functional landscapes. By analyzing many highly divergent homologs, individual steric clashes, which might be important to a particular sequence, become averaged across the orthologs. In addition, the protocol is perhaps simpler than current deep-scanning approaches as the synthesis is multiplexed. More broadly, DropSynth opens entirely new classes of genetic elements to be probed by multiplexed functional assays including proteins and domains, eukaryotic promoter regions, and longer gene regulatory regions and exon/intron junctions. In addition, unlike mutagenic approaches, DropSynth allows for precise designs of libraries to be tested, which allows larger-scale testing of design tools such as Rosetta^5^. The scale and quality of DropSynth libraries can likely be improved further with investment in algorithm design, better polymerases, and larger barcoded bead libraries. We also show that DropSynth can be combined with dial-out PCR^15^, which could be expanded for gene synthesis applications where perfect sequences are paramount. Finally, we envision many potential applications for this approach beyond gene synthesis such as assembling designed gene/pathway combinations and using degenerate oligos to locally explore sequence space around many distant sequences. Importantly, the method requires an initial investment in a barcoded bead set, but then is experimentally tractable using vortexed emulsions, no specialized equipment, with estimated total costs below $2 per gene (Supp. Table 5 & 6).

## Acknowledgements

This work was supported by the funds from the Human Frontier Science Program [LT000068/2016 to C.P.], Netherlands Organisation for Scientific Research Rubicon fellowship [to C.P.], National Science Foundation Graduate Research Fellowship under Grant No. 2016211460 [to A.M.S], National Institutes of Health New Innovator Award [DP2GM114829 to S.K.], Searle Scholars Program [to S.K.], Department of Energy (DE-FC02-02ER63421 to S.K.), UCLA, and Linda and Fred Wudl. We thank George M. Church and Richard Terry for guidance during the early developments.

## Methods

### Design of PPAT library

PPAT orthologs were found by running a PSI-BLAST search with 1 iteration querying the NCBI RefSeq non-redundant protein database using *E. coli* PPAT (NP_418091.1). The resulting set of 11,062 orthologs was further pruned to 10,277 by keeping only those with lengths ranging from 100 to 200 amino acids. T-Coffee (v11) was used to align the sequences and RAxML (v8.2.10) to infer a maximum-likelihood phylogenetic tree (Supp. Fig. 1). A custom Python script trimmed the tree by determining the distance from the root at which the number of nodes equaled the desired amount of orthologs, and subsequently choosing a random descendant leaf for each node at that distance. This reduced the tree around 1,300 ortholog proteins. Each leaf on the pruned tree was then compared to its nearest neighbours to ensure neighbouring sequences differed by at least five amino acids. We then added in orthologs from several model organisms and 34 pathogenic organisms. The final library was dispersed among three libraries of 384 orthologs, with every other leaf on the tree distributed into a different library, for a total of 1,152 orthologs. The final library contained members sourced from 3 Archeal, 9 Eukaryotic, and 1140 Bacterial organisms (of which the top four most represented Bacterial phyla were 414 Firmicutes, 337 Proteobacteria, 64 Actinobacteria and 38 Spirochaetes).

### DropSynth barcode design

We took 2,000 20-mer primers whose design was previously described^25^ and removed those containing NdeI, XhoI, EcoRI, KpnI, NotI, SpeI, BtsI, or BspQI restriction sites. All possible 12-mer subset primers were generated and screened for self-dimers, GC content between 45% and 55%, and melting temperature between 40°C and 42°C. Barcodes were further filtered to ensure a minimum modified Levenshtein distance of 3 between any selected barcodes^26^. The first 384 12-mer barcodes, were used in subsequent oligo designs, with the complementary barcode sequences used to generate the beads.

### Oligo design

Our oligo design protocol is adapted from Eroshenko et al^25^ and summarized in Supp. Fig. 15A. Briefly, protein sequences were assigned a weighted random codon, in order to generate a nucleotide sequence compatible with the restriction sites required (NdeI, KpnI, BtsI, or BspQI). A KpnI restriction site (*GGTACC*) was added on the C-terminal end of the coding sequence, which encodes for a glycine and threonine, before the stop codon. The NdeI restriction site (*CATATG*) on the N-terminal defined the start ATG codon of the ORF. Immediately flanking these restriction sites, 20-mer assembly primer sequences were added, which are used in the emulsion PCA. These sequences were then split into five shorter overlapping fragments^25^, with overlaps optimized to be around 20 bp with a melting temperature between 58°C and 62°C. Sequences which failed to split with these parameters had a new weighted random codon assignment generated, until a codon sequence was found which could be split successfully. BtsI sites were subsequently added on either side of the split sequences, which would release the sequences required for assembly from the bead inside the emulsion droplets, allowing the PCA to proceed. A padding sequence consisting of ATGC repeats was added on the 5’ end ahead of the first BtsI site, with the repeat length such that the final sequence length was 142 nt. Subsequently, an 8-nt Nt.BspQI site, the corresponding 12-mer DropSynth barcode (described above), and another Nt.BspQI site was prepended to the 5’ end of the sequence, with the restriction sites oriented to nick the top strand on the 5’ side of the barcode and the bottom strand on the 3’ side of the barcode. These Nt.BspQI sites facilitate the processing of the barcode region into a single-stranded top-strand overhang. The barcode was common to all five fragments for each gene, such that all fragments required for each gene assembly would be pulled down and localized onto the same beads. Finally a pair of 15-mer amplification primer sequences were added, with each pool of 384 genes (1,920 oligos) having a unique primer pair orthogonal to the other pools. BLAT^27^ was used to screen these primers against the oligo sequences, removing those with homologies over 10 bp. After each of these design steps, we screened for the addition of illegal restriction sites, and modified the sequence respectively if any were found. Three libraries of 384 genes as well as a small test library of 24 genes were ordered as a single pool of 5,880 oligos and synthesized on a microarray by Agilent Technologies.

### DropSynth barcoded beads protocol

The general strategy for creating the DropSynth barcoded beads is shown in Supp. Fig. 15B. Three oligos are used for each DropSynth barcoded microbead, with two of the oligos common to all beads. The anchor oligo attaches to the streptavidin bead surface through a double biotin modification on the 5’ end and has sequences necessary to hybridize with the ligation oligo and part of the barcode oligo. The ligation oligo has a biotin modification on the 3’ end and phosphate group on the 5’ which allows it to ligate to the DropSynth barcode oligo. A different DropSynth barcode oligo is synthesized for each barcode with a common sequence on the 3’ end which can hybridize to the anchor oligo and the reverse-complement of the DropSynth barcode on the 5’ end which can pull down the gene fragments. This approach means only two synthesized oligos (anchor and ligation oligos) contain expensive modifications. Briefly the anchor oligo, ligation oligo, and each barcoded oligo are hybridized, ligated, and phosphorylated with T4 PNK. These are bound to streptavidin coated M270 Dynabeads, washed, and pooled together to form a uniform mixture of all 384 barcoded beads. This protocol can be scaled as necessary given the amount of multiplexing required. The current assembly protocol utilizes 18 uL of the final pooled bead mixture (~3.25E5 beads/uL) for the capture of processed oligos, with the bead barcoding protocol provided producing enough pooled beads to carry out around 210 assemblies in 384-plex.

### DropSynth protocol

DropSynth assembles gene-length fragments through the hybridization of oligos to barcoded microbeads and their resulting amplification. Briefly, individual oligo libraries are PCR-amplified using KAPA HiFi and 15-mer amplification primers. Oligo subpools are then bulk-amplified using the reverse amplification primer and a biotinylated forward amplification primer. After amplification, oligos are nicked using the nicking endonuclease Nt.BspQI, exposing a 12-nt ssDNA “barcode” overhang (Supp. Fig. 16, Supp. Table 3). The short biotinylated fragment that is cleaved following nicking is then removed by binding it to streptavidin M270 Dynabeads in a hot water bath. After a column cleanup, each oligo subpool is mixed with the designed DropSynth barcoded beads and Taq ligase, and annealed overnight from 50°C to 10°C. In this process, all oligos required for each gene assembly are captured when each DropSynth barcode overhang anneals to a corresponding complementary DropSynth barcode on the bead. Captured beads are then mixed with KAPA2G Robust Mastermix, 20-mer forward and reverse assembly primers, BSA, BtsI, and BioRad Droplet Generation Oil. The mixture is immediately vortexed for 3 minutes, allowing for compartmentalization of captured beads in <5 um droplets (Supp. Fig. 17), which are subsequently heated allowing temperature-sensitive BtsI to release the sequences required for assembly from the bead. Droplets from each subpool are then loaded into PCR tubes and thermocycled, allowing PCA to proceed. The PCA products are then recovered by breaking the emulsion with chloroform, purified and re-amplified, providing sufficient assembled DNA for downstream applications.

### Optimization of DropSynth

Significant optimization of the oligo processing and bead capture was required to achieve sufficiently high specificity to allow large multiplexing. Initial attempts to capture fully single-stranded oligos, generated using USER / ƛ exo / DpnII treatment^10^, followed by primer extension of the missing complementary strand, performed poorly for three-oligo assemblies and failed altogether with four-oligo assemblies for all four polymerases tested (Kapa Robust, Kapa HiFi, Pfu Turbo, and Phusion). As an alternative approach, we nicked opposite strands on either side of the BC region with type IIS enzymes, before melting the DropSynth barcode strands apart and removing the unwanted biotinylated strand, leaving a single-stranded overhang along with the rest of the oligo, as shown in Fig. 1A. This eliminated the need for primer extension, and resulted in a 10-fold specificity improvement in tests on 96-plex assemblies of three to six oligos. We further optimized the amount of beads, ligation reaction, fragment capture, the ligase used in the capture step, nicking reaction, and different techniques to purify the emulsion assembly products before re-amplification to achieve an assembly enrichment factor of 10^8^, relative to the probability of a correct assembly by random chance, for a 288-plex five-oligo assembly (Fig. 1B).

### PPAT rescue plasmid and coaD knockout

As PPAT (*coaD*) is an essential gene, we re-engineered plasmid pTKRED28 and to constitutively express bicistronic wild-type (WT) *coaD* gene followed by sfGFP (Supp. Fig. 6A). The WT *coaD* gene from *E. coli* MG1655 was amplified with a strong constitutive promoter (<u>*TTGACG*</u>GCTAGCTCAGTCCTAGG*TACAGT*GCTAGC) and RBS (TACGAGTG<u>*AAAGAGGAGAAA*</u>TACTAG) on the 5’ end, and BamHI site on the 3’ end. This was ligated to a fragment containing a 5’ BamHI site, RiboJ self-splicing element^29^, sfGFP^30^, and a transcriptional terminator to create coaD_sfGFP. pTKRED was digested with BsaI and the larger fragment (8,391 bp) containing the λ-red genes was gel extracted. The coaD_sfGFP DNA fragment was then ligated into the larger pTKRED BsaI fragment to create pTKcoaD. This ligation was transformed into NEB 5-alpha electrocompetent *E. coli* and colonies were sequence verified. The pTKcoaD plasmid expresses PPAT and GFP constitutively while the λ-red recombinase genes are under IPTG induction. The temperature sensitive origin of replication can be used to heat cure the plasmid at 42°C, which can be confirmed through the loss of GFP fluorescence (Supp Fig. 6E).

Knockout of the *coaD* gene in *E. coli* was carried out using standard techniques^28,31^. Briefly pTKcoaD was transformed into both *E. coli* DH10B electrocompetent cells (ThermoFisher Scientific). Individual colonies were chosen and made electrocompetent. These were transformed with a recombination template containing a Kanamycin cassette flanked by homology arms to the regions immediately adjacent to the *coaD* gene. This template was made by first amplifying the Kanamycin cassette from pZS2-123^32^ using primers coaD_KO_KAN_FWD_1 and coaD_KO_KAN_REV_1. The resulting amplicon was purified and further amplified using the primers coaD_KO_KAN_FWD_2 and coaD_KO_KAN_REV_2. The knock-in targeted only the PPAT coding region so as to not interfere with the essential waaA gene immediately upstream of *coaD*. Knock out strains were verified by Sanger sequencing and colony PCR (Supp. Fig. 6C, 6D). We further verified that heat curing of the rescue plasmid suppressed cell growth and characterized the escape frequency.

### pEVBC expression plasmid

The plasmid used to express PPAT orthologs is a derivative of high-copy pUC19 with a pLac-UV5 promoter, NdeI and KpnI restriction sites for cloning, an in-frame stop codon, and a 20-mer random assembly barcode. This was made by first double-digesting pUC19 with AatII + BspQI and gel extracting the larger fragment. A gBlock DNA fragment was synthesized containing the promoter, several restrictions sites, and an in-frame chloramphenicol acetyltransferase before the stop codon. This was ligated into the pUC19 AatII-BspQI backbone fragment to create plasmid pEV_CMR. The plasmid pEV_CMR was double digested with NcoI + KpnI and the long 2,209 bp fragment was gel extracted. Round-the-horn PCR was carried out using 1 ng of the pEV_CMR digest as template, a forward primer pEVBC_FWD with a 5’ biotin and a NdeI site, and a reverse primer pEVBC_REV1 with a 5’ biotin, a 20 N-mer random assembly barcode, and a KpnI site, for 5 cycles. This PCR product was further amplified with outer primers pEVBC_FWD and pEVBC_amp_FWD for 15 cycles. This amplicon was column purified, digested with NdeI + KpnI, treated with rSAP, cleaned up with Streptavidin coated Dynabeads to remove the small fragments, and column purified again to create the vector pEVBC (Supp. Fig. 6B).

### Barcoded PPAT library in pEVBC

Assembled PPAT ortholog genes for each library were digested with NdeI + KpnI and column purified. A ligation was then carried out for each PPAT library using 150 ng of NdeI + KpnI digested pEVBC vector and 100 ng of digested PPAT ortholog genes using 3,000U of T7 ligase in a total volume of 30 uL. This reaction was column purified and concentrated to a volume of 16 uL. NEB 5-alpha electrocompetent *E. coli* cells were then transformed using 3-4 uL of the purified ligation, resulting in over 10 million cfus per transformation. Overnight cultures grown in LB with Carbenicillin were miniprepped, quantified, and an equimolar pool from all three PPAT ortholog libraries was created, henceforth referred to as sample S0.

### Assembly barcode mapping

The assembly barcoded PPAT libraries were sequenced on two Illumina Miseq paired end 600-cycle runs. Each PPAT library was PCR amplified using primers mi3_FWD and mi3_REV_N7## to add p5 sequencing adapters and library indexes. The resulting amplicons were size-selected using gel-extraction and quantified using an Agilent 2200 Tapestation. Samples were then pooled and sequenced on a Miseq using custom primers mi3_R1, mi3_R2, and mi3_index. This resulted in 27,822,356 total reads after merging the runs together. Barcode read counts for S0 were generated by extracting the 20 bp sequence corresponding to the barcode region from the Read 2 sequences and using Starcode^33^ to collapse barcodes within a Levenshtein distance of 1 (Supp. Fig. 3).

Briefly, the subsequent data processing was carried out as follows. All Fastq files had adapters trimmed in bbduk followed by paired-end read merging using bbmerge (from the BBTools package version 36.14). All reads were then concatenated and piped into a custom python script which generated a consensus nucleotide sequence for each barcode. The script works as follows. First, we split reads into the 20 nt assembly barcode and the corresponding variant, and generate dictionary that maps every assembly barcode to a list of variants associated to it. To eliminate assembly barcodes that are associated with two different variants, we calculate the pairwise Levenshtein distance of every variant associated with a given assembly barcode. If a certain percentage of these assembly barcodes (5%) are greater than a distance cutoff (10) then we consider the assembly barcode contaminated and drop it from further analysis. Finally, we generate a consensus sequence by taking the majority basecall at every position. Mapped consensus sequences were then translated until the first stop codon and sequences perfectly matched to any designed orthologs were annotated.

Analysis of the number of reads per assembly barcode as a function of dilution revealed a small number of assembly barcodes with very high number of reads, as many as 300,000 by the fourth dilution, attributed to the emergence of adaptive mutations conferring a growth advantage at 42°C, which occur stochastically. We also deduce from the lack of GFP positive colonies in the plates at various steps in the dilution that these adaptive mutations did not occur in cells still harbouring the rescue plasmid. A total of 18 barcodes from serial dilution replicate A and 16 barcodes from replicate B were removed from further analysis.

Mutant ortholog sequences were annotated by first aligning the consensus nucleotide sequence for each barcode against the 1,152 designed PPAT orthologs using bbmap. The resulting SAM file was parsed to extract the closest alignment match. A pairwise alignment of the amino acid sequence was carried out for each mapped barcode sequence (until the first stop codon) against its best PPAT ortholog alignment match. Mutants within a distance of 5 amino acids from the designed sequence had their individual a.a. mutations annotated for further analysis downstream.

### Complementation assay

The complementation of synthesized orthologs was carried out using a serial batch culture. After ligation into pEVBC, orthologs from all three libraries were pooled together to create sample S0. Supercoiled S0 plasmid was then electroporated into electrocompetent *E. coli* DH10β *ΔcoaD* pTKcoaD. The serial batch cultures, consisting of two biological replicates, were initiated by making 10 transformations using 1 ng of S0 plasmid into 40 uL of cells and recovered at 30°C in 1mL SOC + 1 mM IPTG for 1 hour. For each replicate, 5 transformations were pooled together and used to seed a fresh culture with between 7 million and 17 million cfus. Cells were grown in 1 L LB media supplemented with Kanamycin + Carbenicillin + 0.05 mM IPTG and grown to saturation at 42°C (8-10 generations) between each bottleneck. Cells were propagated through 3 bottlenecks for a total of 4 samples for each replicate, with 1000x dilutions at each bottleneck. DNA was miniprepped from each sample and cells were plated to ensure proper curing of the rescue plasmid, by screening for GFP+ colonies.

The barcodes from each of the 8 complementation samples were amplified using primers mi4_FWD and mi4_REV_N7## to add sequencing adapters and library indexes. The resulting 294 bp amplicon was size-selected using gel-extraction, purified, pooled, and loaded onto a Hiseq 2000 single-end 50 cycle run using custom sequencing primers mi4_R1 and mi4_index, resulting in 138 million total reads. The barcodes for each sample were clustered using Starcode^33^ to collapse barcodes within a Levenshtein distance of 1 (Supp. Table 1).

### Complementation data analysis

In order to reduce noise in calculating the fitness change we pruned the barcodes leaving only those with at least 10 reads in S0 or at some point in the serial dilution. This reduced the total number of unique barcodes from 7,038,274 to 627,302. Further data analysis was carried out in R, with visualisations using ggplot2 and ggtree^34^. We calculated fitness scores for each mapped sequence with at least one barcode. First, the read counts at each dilution were normalized based on the total sequencing depth of the sample relative to S0. The log2 fold change between each dilution and sample S0 was then determined for each barcode using

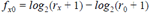

where *r*_*x*_ is the number of normalized reads in the corresponding dilution. We then took the median value (to minimise effects of outliers) of the log2 fold change over all of the dilutions to determine the fitness for that barcode

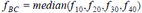

The median fitness for each barcode representing a sequence was determined for each replicate (A and B) individually

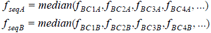

We then selected only those sequences represented in both replicates and took the median replicate fitness as the final fitness value

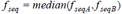

### Assembly Retrieval by Dialout Amplification

The presence of a unique barcode on each assembly allows us to retrieve them from the library using PCR amplification^13,15^. We attempted to amplify 48 unique orthologs and 12 gain-of function mutants. As a positive control we also amplified the wild-type *E. coli* coaD gene from the pTKcoaD rescue plasmid. The designed primers flanked each construct, with reverse primers annealing to each gene-specific barcode. We observed correct size amplification products for 59 of 60, with 18 of these using lower complexity post complementation selection libraries as template, while the rest used the high-complexity sample S0. Individual amplicons were then gel-extracted, restriction digested with KpnI-HF and NdeI, ligated into empty pEVBC backbone, and transformed into chemically competent NEB DH5alpha *E. coli* cells. Colonies were verified via colony PCR and Sanger sequencing, and validated colonies were re-inoculated overnight and miniprepped. We successfully sequence-verified 43 of the 59 constructs (37 orthologs and 6 gain-of-function mutants), in addition to the WT coaD gene (Supp. Table 2).

### Growth Rate Analysis of Dialed-out Orthologs and Gain-of-function Mutants

Following successful dialout PCR and re-cloning, we transformed 1 ng of each construct in pEVBC into 7 uL of electrocompetent *coaD* knockout cells. We analyzed the presence of growth by counting dilution Carb + Kan plates at both 30°C and 42°C (Supp. Table 3, Supp. Fig. 10A). Four constructs had no colonies on the 42°C plates, of which two were low-fitness orthologs, one was a gain-of function mutant with only one barcode (false positive), and another construct KOS35328 had good fitness (1.88) in the pooled assay determined using 25 barcodes. The lack of colonies for KOS35328 requires further investigation, and may be a transformation error. Six constructs had low colony counts on both plates, of which five correspond to low-fitness orthologs (Supp. Fig. 10A). We noticed a trend in which orthologs with enhanced fitness in the pooled complementation assay gave rise to greater numbers of colonies on the 42°C dilution plates. Furthermore, we also noticed that orthologs with enhanced fitness in the pooled complementation assay typically gave rise to 42°C colonies that appeared larger than their corresponding 30°C colonies (Supp. Figure 12A). Of the constructs with at least 10 colonies on the 42°C plates, we picked 3 colonies per ortholog and re-inoculated them in 1 mL LB + Carb + Kan and grew overnight at 42°C. 2 uL of saturated culture was then diluted in 98 uL of LB + Carb + Kan in wells of a 96-well plate and loaded into a Tecan M1000 Plate Reader for 12 hours at 42°C. OD600 values, taken at 30-minute intervals, were measured at 9 points within each well and averaged. Resultant growth curves were plotted for all colonies and averaged on the construct level. Maximum slopes of each growth curve were calculated and plotted against fitness scores determined from the complementation assay (Supp. Fig. 10B). A strong correlation (Spearman r_s_ = 0.86) was observed comparing ortholog growth rate to fitness, validating our assay and analysis pipeline. Examining the residual errors of the fit of growth rate to fitness we observe that constructs with fewer barcodes tend to have larger errors (Supp. Fig. 10C) which agrees with the reproducibility of the fitness value among replicates as a function of the number of barcodes (Supp. Fig. 7B).

